# Mentally transformed representations in memory are linked to their originals

**DOI:** 10.64898/2026.09.10.750562

**Authors:** Nursena Ataseven, Șahcan Özdemir, Elkan G. Akyürek, Daniel Schneider, Wouter Kruijne

## Abstract

Working memory is an active workspace for manipulating encoded information. Neural decoding studies show that following mental transformation, the encoded original remains represented alongside the transformation product despite offering little to no functional merit. To test why, we employed multivariate EEG-decoding while participants mentally rotated a memorized orientation grating, encoded an additional grating, and were finally retro-cued whether the rotation product or the additional item would be probed. This tested whether retention of the original reflected (1) perceptual encoding, (2) deep encoding and persistent storage, or (3) a link to the transformation product. The original was decodable after rotation, and encoding an additional item, suggesting its retention was not merely perceptual. Next, contingent on the retrocue, the original and rotation product were retained or dropped together: the original remained decodable only when the rotation product was selected. Therefore, the original did not persist because it was deeply encoded; its retention depended on attentional selection of the transformation product. Yet, we found no evidence that retaining the original was beneficial. Our results show that mental transformation links the transformation product to its source representation, reflecting a dependent representational structure. We speculate that this structure may afford behavioral flexibility in dynamic environments.

## Introduction

The dynamic world that we live in requires a flexible mental workspace to predict the future states of our surroundings to guide behavior. For example, when driving, perceptual information encoded from the rear window has to be mentally transformed to predict the upcoming locations of the approaching cars when gazing forward. Working memory (WM) has been thought to provide such a workspace that not only allows temporarily maintaining information, but also supports its manipulation in the absence of sensory input (Baddeley, 2003; Logie, 2003; Logie & Della Sala, 2003; Postle, 2015; Smith & Jonides, 1999). Such manipulations allow us to transform the maintained representations according to task demands, enabling flexible behavior in ever-changing environments.

It has been suggested that sensory encoding and retention of imagined or transformed internal representations rely on overlapping networks (Albers et al., 2013; Roelfsema & De Lange, 2016; but see also Iamshchinina et al., 2021). Such overlap makes it difficult to determine the representational nature of transformed internal representations. A transformed representation could arise from updating and eventually replacing the originally encoded sensory trace, or from constructing an additional representation while the original trace remains available. Evidence supporting the latter hypothesis has come from previous studies that have investigated the neural representations after mental transformation using a mental rotation paradigm. These studies show that the originally encoded representations remain available after a transformation product has been generated, despite the original representation being explicitly declared task-irrelevant (Christophel et al., 2015; Kandemir et al., 2024). Neural evidence from trial-wise decoding indicates that the representations are held concurrently, rather than competing with each other. Such persistent and concurrent maintenance of an irrelevant representation is surprising: studies on WM representations in retrospective cueing paradigms typically show that as soon as information is deemed task-irrelevant, it can be purged from WM quickly and efficiently (Griffin & Nobre, 2003; Landman et al., 2003; Oberauer, 2001; Schneider et al., 2016; Souza et al., 2014; Wolff et al., 2017; Zanto & Gazzaley, 2009).

To investigate why originally encoded representations were found to persist after they became task-irrelevant, we outline three preregistered (<u>osf.io/f6w2s</u>) hypotheses. The first hypothesis is that the original representation is retained as a consequence or by-effect of perception or WM encoding, and that it’ll be represented only until new perceptual input is encoded. The second hypothesis is that a mental transformation requires an original representation to be deeply encoded into WM, and that its persistent retention is a consequence of this extensive encoding process. Under this account, the original representation would persist throughout the task, regardless of the task-relevance of the original representation or the transformation product. The third hypothesis is that the transformation product may be maintained in a higher-order task representation generated from the originally encoded representation, such that the original representation is retained in a manner that is functionally linked to the transformation product. In such a case, the original item is only retained when the transformation product is task-relevant, and is dropped when the transformation product is task-irrelevant.

In the present study, we tested these predictions using a WM task with mental rotation. We recorded EEG while participants memorized and mentally rotated an orientation grating, with the outcome of this transformation referred to as the transformation product. Participants then memorized an additional orientation grating. A retrocue subsequently indicated whether the transformation product or the additional grating was to be probed, allowing us to direct attention towards or away from the transformation product, and observe the consequences for the originally encoded representation. To track WM representations, we presented an impulse stimulus during the retention intervals and analyzed the resulting EEG response (Wolff et al., 2015, 2017). If the original representation disappeared after the additional item was encoded, regardless of whether the transformation product was later selected or deselected by the retrocue, then the first hypothesis, which considers the maintenance of the original item is a consequence of perceptual encoding, would be supported. If the original representation persisted throughout the task regardless of the retrocue, then the second hypothesis, which considers the maintenance of the original as a byproduct of deep-encoding, would be supported. Finally, if the original representation persisted when the transformation product was selected by attention, but disappeared when the transformation product was deselected in favor of the additional item, then the third hypothesis, proposing linked-representations for the original and the transformation product, would be supported.

## Results

Participants performed a delayed comparison task during which they memorized and mentally transformed an orientation grating (Figure 1A). On each trial, they first memorized an orientation grating, referred to as the original item. Subsequently, they rotated this item mentally (+-30°, 60°, 90°). The resulting transformed representation is referred to as the rotation product. In these trials, the participants were informed that only the rotation product was task-relevant and that the original item was never probed. On 25% of trials, the instruction was 0°, indicating that no mental rotation was required. The target in these trials is referred to as the no-rotation original item. Next, an additional orientation grating was displayed to be memorized, which is referred to as the additional item. This was then followed by a retrocue indicating whether the rotation product (retrocue “1”; or the original item following a no-rotation instruction) or the additional item (retrocue “2”) would be probed. Finally, the participants judged whether the retrocued orientation was oriented clockwise (CW) or counterclockwise (CCW) compared to the probe orientation presented on the screen.

**Figure 1.**
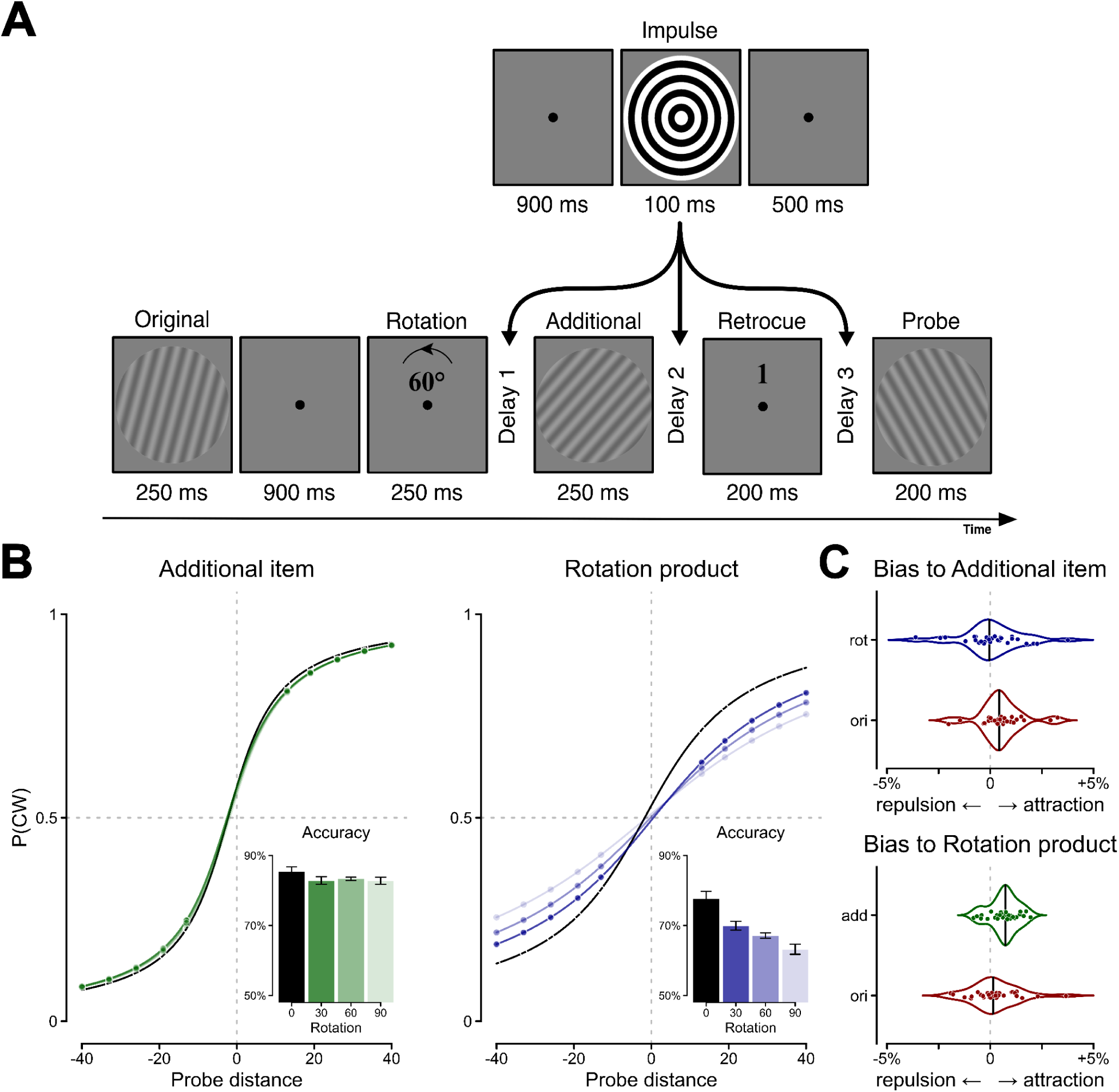
Experimental procedure and behavioral results. **(A)** Illustration of a trial. Participants first memorized the orientation of a grating, referred to as the original, and later rotated it. Subsequently they memorized the orientation of an additional grating. A retrocue either indicated that the rotation product (“1”) or the additional item (“2”) was to be tested. The participants judged whether the probe grating was CW or CCW relative to the cued item. An impulse was shown in each of the delay intervals to assess WM representations during maintenance. **(B)** Probability of giving a CW response as a function of probe distance and absolute rotation magnitude in trials where the additional item (left panel) or the rotation product (right panel) was cued. The psychometric curves reflect the binomial regression fit evaluated at each absolute rotation magnitude. Dots on the curves indicate the tested probe distances. The inset plots depict the mean accuracy for each absolute rotation magnitude with error bars reflecting within-subjects confidence intervals per magnitude. **(C)** Bias to the additional item (upper panel) or the rotation product (lower-panel) by the other two items. The plots reflect the predicted percentage increase in CW responses associated with a 10-degree increase in CW probe distance to either bias source item.

### Behavioral results

We modeled CW responses as a psychometric function of probe distance, using hierarchical binomial regression, and assessed effects on precision as the interaction terms modulating the slope of this function. The best model indicated that precision was modulated by which item was probed (retrocue) and by whether mental rotation took place on the trial. Precision was higher for the additional item than the rotation product (*χ²*(2) = 1187.49, *p* < 0.001). In addition, precision was higher on trials with no mental rotation than trials with nonzero rotation magnitude (*χ²*(2) = 234.98, *p* < 0.001). Tests for a three-way interaction between these effects provided marginal evidence for slightly underadditive benefits of cueing the additional item and performing no mental rotation (*χ²*(2) = 5.4, *p* = 0.068).

When evaluating only the trials where the additional item was cued, we again found that no-rotation trials showed slightly better precision (*χ²*(2)=11.83, *p* = 0.003). The absolute magnitude of rotation, however, did not affect precision on these trials (*χ²*(2)=1.16, *p* = 0.560). Conversely, trials where the rotation product was cued showed both a benefit for no-rotation trials (*χ²*(2) = 30.08, *p* < 0.001), and a linear effect of the magnitude of mental rotation: larger mental rotation lead to decreased precision (*χ²*(2) = 56.89, *p* < 0.001).

Figure 1B summarizes these results: across conditions, participants performed well above chance, but performed better when the additional item was probed than the rotation product. On trials where the rotation product was probed, performance decreased as a consequence of mental rotation. In addition, we found that performance was better on both items when no rotation took place on the trial, presumably because this requires less mental effort overall. As trials with no rotation are not comparable to the trials with mental rotation and also not of interest to our research question, they are excluded from all subsequent analyses.

To better understand the functional properties of the original representation following mental rotation, we assessed to what extent it affected behavior. To this end, we computed the distance between the probe stimulus and each of the three items: the original, the rotation product, and the additional item. We split the data by retrocue, and assessed to what extent the distances between the probe and other items biased responses to the cued item (Figure 1C). We quantified these biases by computing the predicted increase in percentage of CW responses across all probe distances, given a 10 degree increase in the signed distance between the probe and the other item.

On trials where the additional item was probed, responses were slightly attracted by the task-irrelevant original item (*χ²*(1)=5.18, *p* = 0.023, *M_bias_* = +0.52%). The uncued rotation product, however, did not seem to affect behavior (*χ²*(1)=0.58, *p* = 0.448, *M_bias_* = +0.09%). Conversely, trials where the rotation product was probed showed no bias from the task-irrelevant original item (*χ²*(1) = 0.88, *p* = 0.348, *M_bias_* = +0.23%), yet there was an attractive bias from the uncued additional item (*χ²*(1) = 31.77, *p* < 0.001, *M_bia_*_s_ = +0.66%).

Together, these results indicate that while perception and encoding of the original and additional item seemingly interfered with WM representations, the original item representation did not modulate responses to the rotation product. This suggests that the persistent representation of the original item, which will be reported in the next section, does not stem from a failure to comply with rotation instructions. In addition, it suggests that the original item is not used as a ‘fallback’ item to guide responses.

### EEG decoding of rotation trials

In the subsequent analysis, we decoded from trials with a mental rotation instruction. We report the average decoding results within the 0–300 ms time window following each impulse onset during the delay intervals, as well as time-resolved decoding results across the whole epoch. During Delay 1 (see Figure 1A), the rotation product was successfully decodable from the impulse display (Figure 2A, whole-epoch significant cluster: 4–432 ms, *p* < .001; time-window average: BF_10_ = 7.74, *p* = .005), suggesting that the participants performed mental rotation successfully. Replicating previous findings (Christophel et al., 2017; Kandemir et al., 2024), the original item was also decodable after mental rotation during Delay 1 (whole-epoch significant cluster: 28–412 ms, *p* < .001; time-window average: BF_10_ = 89.52, *p* < .001). Therefore, even though the original item was task-irrelevant at this stage, it was still kept in memory alongside the task-relevant rotation product.

**Figure 2.**
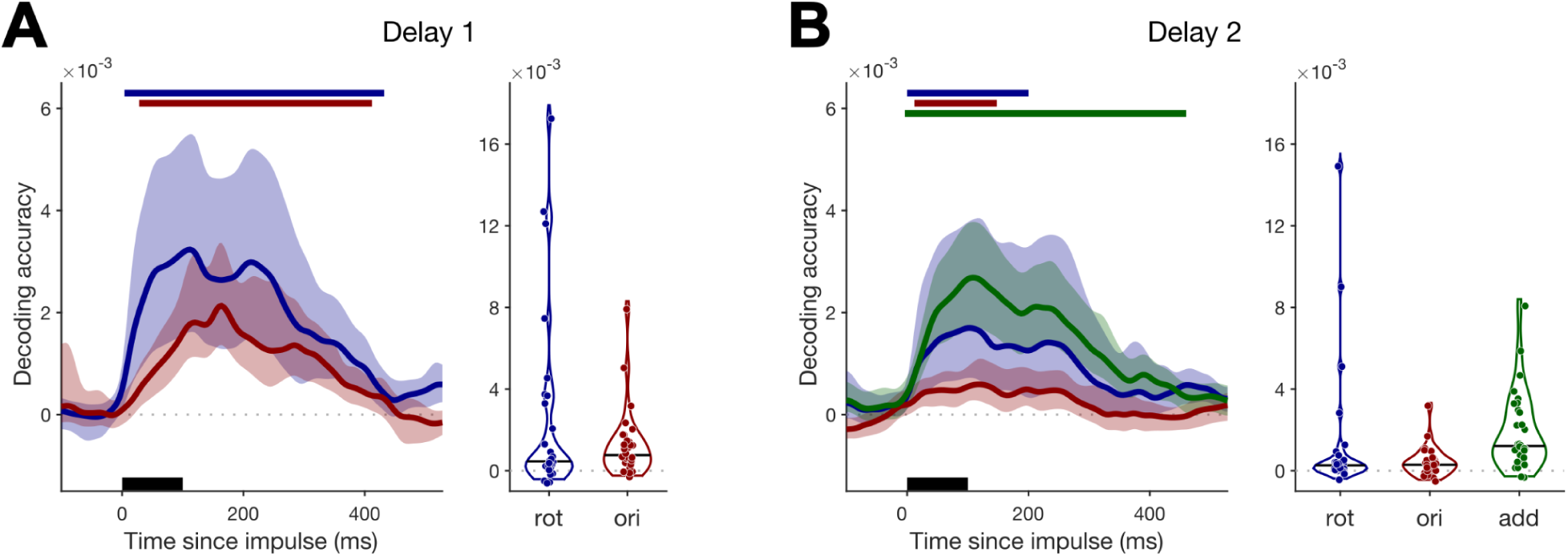
Decoding results from Delay 1 (A) and Delay 2 (B). The time-course plots are time-locked to the impulse onset in Delay 1 and 2 marked by the black lines over the x-axes. The solid colored lines show the mean decoding accuracy for the items, with the shaded regions reflecting the 95% confidence intervals. Top horizontal lines mark clusters with statistically significant decoding (p < 0.05, one-sided) for each item. Violin plots show decoding accuracy averaged across the 0-300 ms time window. Dots reflect individual participant means and the black horizontal line in the violins reflects the median.

During Delay 2, the additional item was decodable from the impulse (Figure 2B, whole-epoch significant cluster: −4–460 ms, *p* < .001; time-window average: BF_10_ = 3076.9, *p* < .001)^1^. For the rotation product, we found marginal evidence for successful decoding in our time-window analyses (BF_10_ = 1.48, *p* = 0.038). Within the time window, we found a significant cluster (whole-epoch significant cluster: 0–196 ms, *p* = .001). Importantly, the original item was also decodable following Impulse 2 (whole-epoch significant cluster: 12–148 ms, *p* = .003; time-window average: BF_10_ = 12.37, *p* = .003). This means that despite encoding an additional item, the original item was still represented, suggesting sustained maintenance.

During Delay 3, after the retrocue, we tested the effects of attentional selection on the neural representation of the three items. First, we considered trials in which the additional item was cued. During the delay, the additional item was decodable (Figure 3A, whole-epoch significant cluster: 20–424 ms, *p* < .001; time-window average: BF_10_ = 362.66, *p* < .001), whereas both the original (time-window average: BF_10_ = 0.19, *p* = .906) and the rotation product (time-window average: BF_10_ = 0.34, *p* = .278) were not. This shows attentional selection of the additional item, whereas the task-irrelevant rotation product and original item were discarded from memory.

**Figure 3.**
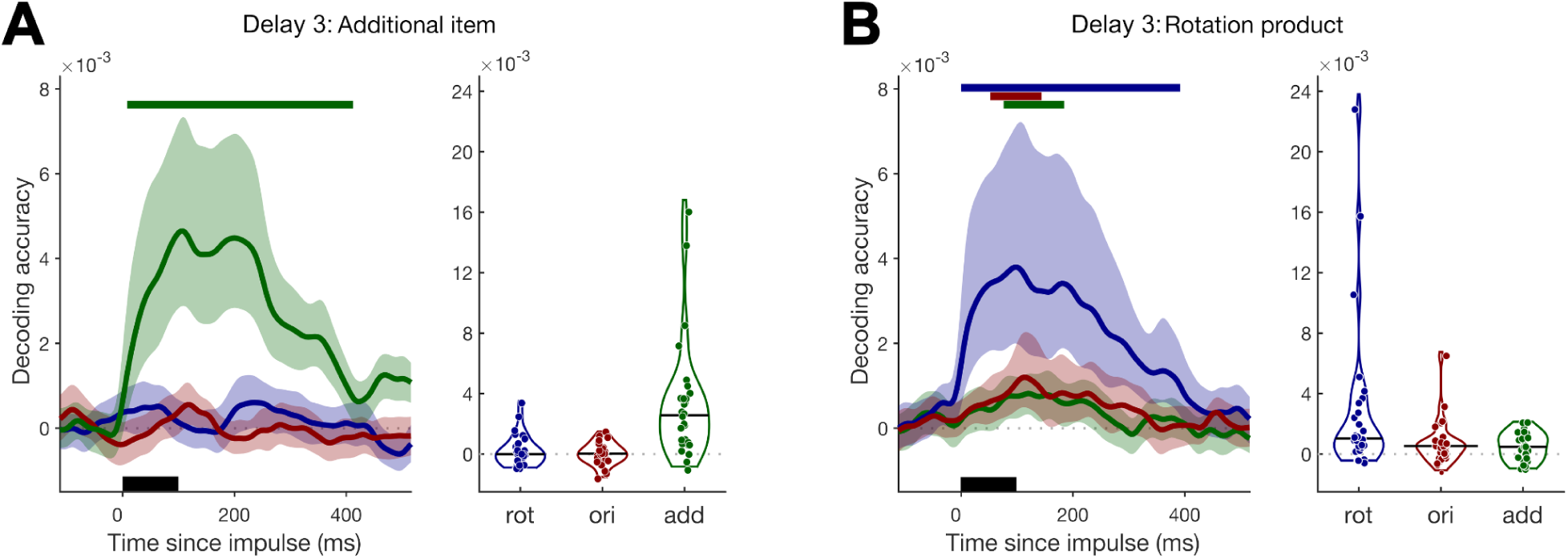
Decoding results from Delay 3 on trials where the additional item (A) or rotation product (B) was cued. The time-course plots are time-locked to the impulse onset in Delay 3 marked by the black lines over the x-axes. The solid colored lines show the mean decoding accuracy for the items with the shaded regions reflecting the 95% confidence intervals. Top horizontal lines mark clusters with statistically significant decoding (p < 0.05, one-sided) for each item. Violin plots show decoding accuracy averaged across the 0-300 ms time window. Dots reflect individual participant means and the black horizontal line in the violins reflects the median.

During trials in which the rotation product was cued, expectedly, the rotation product could be decoded from impulse (Figure 3B, whole-epoch significant cluster: 8–404 ms, *p* < .001; time-window average: BF_10_ = 8.58, *p* = .005). Crucially, the original item could also be decoded in these trials, alongside the rotation product (whole-epoch significant cluster: 64–156 ms, *p* = .023; time-window average: BF_10_ = 3.43, *p* = .014), suggesting that it was attentionally selected along with the rotation product, which was the only one to be probed. Meanwhile, the additional item was also decodable (whole-epoch significant cluster: 88–184 ms, *p* = .012; time-window average: BF_10_ = 7.92, *p* = .005), which may have resulted from the fact that it was recently encoded, rather than task-relevance per se. These results suggest that the original item’s representation is not simply persistent, but is contingent on the attentional selection of the rotation product.

### Relating trialwise EEG decoding to behavior

The decoding results presented thus far suggest that persistent maintenance of the original item is not reflecting passive persistence, but is conditional on the relevance of the rotation product. This could either mean that these item representations are linked in WM, or it could mean that these results reflect a mixture of trials with different strategies, where participants rely on the rotation product on some trials, and the original item on other trials. To address this difference, we exclusively looked at trials where the rotation product was cued, and assessed how trial-wise decoding scores z-scored per participant in Delay 3 related to each other and to behavior.

Interestingly, we found that decoding scores for the original item and the rotation product were positively related at the trial-level: Linear mixed-effect models of rotation product decoding scores showed that the preferred model included a predictor reflecting original item decoding (*χ²*(2) = 55.83, *p* <0.001). Figure 4A illustrates this relation as the average rotation product decoding score as a function of the original item decoding score, split into three bins per participant. As the figure illustrates, slopes across participants were consistently positive (*β* values ranging from 0.102 to 0.234). If the participants had employed a mixture of strategies, one would expect decoding scores to have a negative relationship. Given the positive relationship, these results provide further evidence that the representations of the original item and rotation product are truly linked.

**Figure 4.**
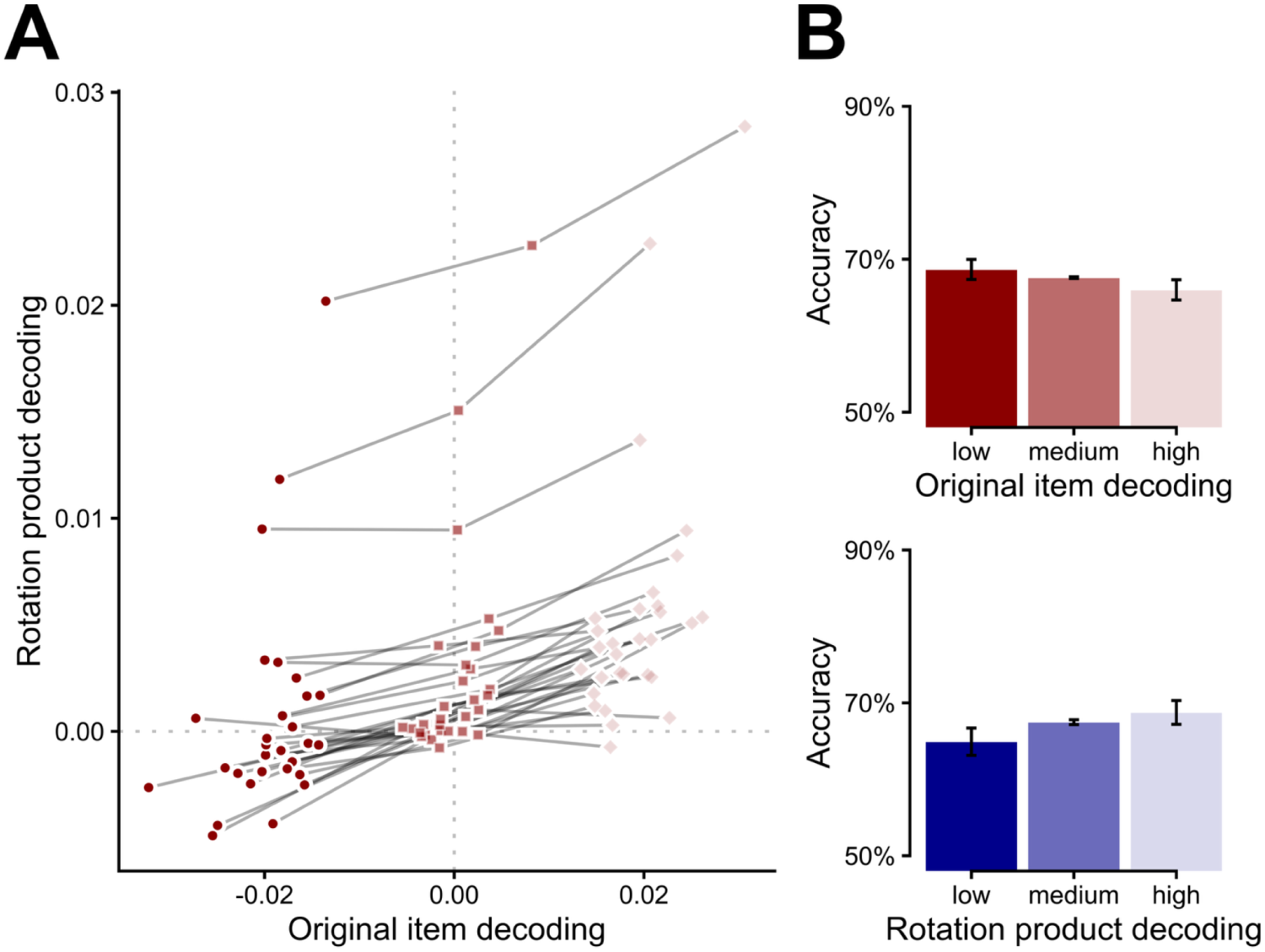
Relationship between trialwise decoding and behavior. **(A)** Trial-level association between z-scored decoding accuracy of the rotation product and the original item. Mean rotation product decoding scores are shown as a function of original item decoding scores, split into three within-participant bins. **(B)** Relationship between behavioral accuracy and z-scored decoding strength of the original item (upper panel) and rotation product (lower panel), with error bars reflecting within-subjects confidence intervals per bin.

To gauge whether relative decoding of either item affected behavior, we first computed the z-scored decoding strength separately per participant and item. Then, we evaluated statistical support for including it as predictor terms in our previously established best model of behavior. We found evidence that z-scored rotation product decoding modulated behavioral precision (*χ²*(2) = 7.91, *p* = 0.019), as did original item decoding (*χ²*(2) = 9.28, *p* = 0.010), without evidence for an interaction between these terms (*χ²*(2) = 0.81, *p* = 0.667). Interestingly though, the two decoding score terms had opposing effects on behavioral performance, despite their positive relationship: Having relatively higher rotation product decoding improved participants’ performance, whereas a higher decoding score for the original item decreased performance (see Figure 4B). These results show that while trials with strong decoding for the rotation product were more likely to have strong decoding for the original item, the residual variance between these two variables reveals opposing marginal effects on behavior.

We next evaluated whether any of these effects would also be present for the decoding score of the uncued additional item. We found that decoding scores of the additional item representation did not correlate with rotation product decoding (*χ²*(1) = 0.02, *p* = 0.895, *β*’s from −0.035 to 0.005), which suggests that the relation between decoding scores for the original item and rotation product does not merely reflect trialwise fluctuations in signal quality. When analyzing behavior, the z-scored decoding scores of the additional item also seemed to have no bearing on behavioral precision (*χ²*(2) = 1.07, *p* = 0.584). Of note, we additionally investigated whether the additional item decoding score was perhaps related to its bias effect on behavior, but again found no support for this (*χ²*(2) = 0.69, *p* = 0.709). So, while original item and rotation product decoding scores in this epoch modulated precision, the additional item exerted a bias on behavior that was not related to its decoding score. This fits with our earlier suggestion that the additional item exerted a bias on behavior during its perception, rather than during maintenance in WM.

## Discussion

Our data reveal that internally generated representations produced through mental transformation remain linked to their initially encoded originals in WM. Specifically, attention operated on both simultaneously: when the transformation product was cued, the original representation continued to be maintained, whereas when the transformation product was not cued, both representations were not decodable. Moreover, the trial-wise decoding strengths of the original item and the transformation product were correlated, yet their marginal effects on behavior were in opposite directions: performance increased when the transformation product was more decodable, while it decreased when the original item was more decodable. Taken together, the results suggest that persistent original item decoding following mental transformation does not reflect trials in which transformation had failed, nor does it reflect that the original is used as a backup or fallback item. Rather, following mental transformation, the original item retains a functional link to the transformation product, such that attentional selection of one representation extends to the other.

In WM paradigms during which participants are asked to memorize and later report an item, task-irrelevant information has been shown to be discarded from WM quickly (Griffin & Nobre, 2003; Landman et al., 2003; Oberauer, 2001; Souza et al., 2014; Schneider et al., 2016; Wolff et al., 2017; Zanto & Gazzaley, 2009). Considering that mental transformations are cognitively effortful (Searle & Hamm, 2017; Shepard & Metzler, 1971; Wexler et al., 1998), discarding task-irrelevant information would allow more efficient usage of cognitive resources. This may appear in contrast to the present results. Yet, our findings are in line with the previous research on mental transformation (Christophel et al., 2017; Kandemir et al., 2024), where continued maintenance of the original item was also observed, even when it was similarly task-irrelevant. In the present study, the maintenance of the original item persisted after mental transformation and after encoding an additional item. These results suggest that the neural trace of the original item is not merely due to extensive encoding of the original item, nor the result of perceptual encoding. Instead it appears to be genuinely retained in memory alongside the transformation product, despite it being task-irrelevant. We observed that the original item seemed to be selected together with the transformation product; it was decodable when the transformation product was cued, but not decodable when it was uncued.

This persistence of the original item may reflect a dissociation between WM storage for representations used in mental transformation and merely visually encoded representations. In attentional cueing paradigms (Griffin & Nobre, 2003; Landman et al., 2003; Oberauer, 2001; Souza et al., 2014; Schneider et al., 2016; Wolff et al., 2017; Zanto & Gazzaley, 2009), the task-irrelevant item is often independent of the selected item, whereas in mental transformation paradigms the task-irrelevant original item serves as the source of the subsequently generated item. These observations raise the question whether maintaining the original item representation serves a particular functional purpose. Our data seem to contradict this: We showed that stronger trialwise original item decoding was associated with lower behavioral performance and responses to the transformation product were unaffected by the original item.

One possible answer is that the neural trace for the original item persists not because it serves a direct functional purpose but because of its representational relationship to the transformation product. The transformation product could constitute an abstracted and transformed representation that is hierarchically nested under the originally encoded representation. Ample evidence suggests that WM is hierarchically organized (Christophel et al., 2017; D’Esposito & Postle, 2015; Fuster, 1997), with lower-level representations being integrated into higher-order chunks. WM representations have been proposed to exist in multiple representational formats, ranging from sensory-like to progressively more abstract and transformed forms (Christophel et al., 2017). Accordingly, the transformation product may be maintained as an efficient higher-order abstraction of the original item, remaining representationally linked to it rather than being encoded as an entirely independent memory trace. Considering that hierarchical chunking is thought to help WM overcome its capacity limitations, a similar organization of the transformation product, nested under the original item, would be plausible, particularly because encoding an additional item in our study did not appear to eliminate the maintenance of the original item. Future research could investigate whether indeed the transformation product constitutes a higher-order abstraction of the original item, maintained in more anterior regions of the cortical hierarchy whereas the original representation remains in the posterior regions in a sensory-like format. Such a separation across hierarchical layers could also explain why Kandemir et al. (2024) did not observe generalizability in the neural code of the transformation product and the original representation.

Of note, while the original item seems to have negatively impacted performance in our task, retaining a representation of the original after mental transformation may actually serve a functional purpose in more naturalistic settings. In the real world, mental transformations may typically serve to simulate how an uncertain situation is likely to develop, based on information retained in WM and dynamically changing incoming sensory information (Moulton & Kosslyn, 2009). For instance, if a car disappears from view in the rear-view mirror, the latest available information about its trajectory can be used to generate a prediction regarding its subsequent location. In such settings, retaining the original representations alongside the transformation product could support cognitive flexibility in case circumstances change (e.g., a sudden change in traffic density). That is, recomputing the mental transformation of the original with different parameters may be more efficient and more accurate than updating a previous transformation product. Thus, although the present task with a single, isolated transformation showed a behavioral detriment of retaining the original item, doing so could also carry a potential benefit of being able to adapt transformations to fallible circumstances, by recomputing them from their originals if needed (see Kandemir et al., 2024, for a similar suggestion).

Another potential function for maintaining the original item alongside the transformation product may be that these are used to build associations that help perform future mental transformation in a more automated manner. In various models of the interplay between WM and reinforcement learning (O’Reilly & Frank, 2006; Ott & Nieder, 2019; Todd et al., 2008; Yoo & Collins, 2022), relevant stimuli encountered while solving a cognitive task leave traces in memory that persist until the end of the trial where reward may be delivered. Having such traces available at the moment of reinforcement allows one to associate these stimuli to the reward and supports temporal difference learning. Of note, attentional selection plays a critical role in shaping these memory traces, such that representations held in memory when reinforcement arrives allow for credit assignment to relevant stimuli and actions while ignoring stimuli deemed irrelevant (Kruijne et al., 2021; Rmus et al., 2021; Rombouts et al., 2015). In the context of the present task, the persistent maintenance of the original item, and its selection being contingent on the task-relevance of the rotation product, could be an expression of a learning process that serves to gradually automate mental rotation skills.

Our findings reveal that mentally generated representations remain dependent on their initially encoded originals, being attentionally (de-)selected together. More broadly, rather than constructing and managing independent representations, the brain may generate transformed representations that are anchored to their respective originals when creating its inner world. The inner world is thus created by representations derived from the external world, while the representations in the inner world remain loyal to the outer world. This mechanism may further reflect an efficient storage mechanism that mitigates the capacity limitations of WM through generating and manipulating novel mental representations without the full cost of maintaining entirely independent representations.

## Materials and Methods

### Preregistration

Hypotheses, data collection^2^, exclusion criteria and analyses of this study were preregistered on an Open Science Framework page (<u>osf.io/f6w2s</u>). All EEG decoding analyses were preregistered while behavioral analysis and the analysis for linking the trialwise EEG decoding to behavior were not.

### Participants

Forty volunteers (26 women and 14 men, mean age: 24.5, range: 18-33) participated in the experiment in exchange for course credit or monetary compensation. Ten participants were excluded from data analysis; four due to low task accuracy (below 60%) and six for not completing the experiment. Participants were informed about the experimental task, EEG data collection procedure, and data sharing procedures, and provided written informed consent prior to experiment. This experiment was conducted in accordance with the Declaration of Helsinki and was approved by the Leibniz Research Center for Working Environment and Human Factors Ethics Committee.

### Apparatus and stimuli

The experiment took place while participants sat in a dimly lit soundproof experimental chamber. The instructions and experimental task were presented on a 19-inch CRT monitor screen with 100 Hz refresh rate and 1,280 by 1,024 pixels resolution from an 80 cm viewing distance. The script for the experimental task was coded and run using MATLAB (R2024a) with Psychtoolbox (Version 3.0.19). Behavioral responses were collected via a keyboard with an USB interface.

As depicted in Figure 1A, the background of the experimental task was set to gray (RGB = [128, 128, 128]). Memory items were circular patches containing sine wave gratings with a frequency of 0.65 cycles/° of visual angle. These items were presented centrally with 20% contrast and a diameter of 6.69° while their phase was randomized across trials. Each memory item was presented in one of six preselected orientations (15°, 45°, 75°, 105°, 135°, and 165°). The rotation instruction consisted of an arc from the top of a circle sweeping from −60° to +60° with an arrowhead positioned at its top center indicating the rotation direction and a numeric rotation magnitude (0°, 30°, 60°, and 90°) placed below. The instruction black (RGB = [0, 0, 0]).

In this display, a numeric rotation magnitude (0°, 30°, 60°, and 90°) that was 2.45° tall was positioned 1.12° above the display center. A circular arc sweeping from −60° to +60° was displayed with 4.63° x 1.34° and was positioned at 2.23° above the center of the screen such that it aligned with the top of the numeric magnitude indication. An arrowhead of 1.28° x 1.11° of visual angle was positioned at the upper center of the arc to indicate the direction of the rotation; a rightward arrowhead for CW rotations, a leftward one for CCW, and no arrowhead for 0° rotation. The impulse stimuli each consisted of a bullseye pattern with the same size and spatial frequency of the memory items but with sharp black-white contrast transitions. The retrocue was a black numeric display (1 or 2) of 2.45° and was positioned at the center of the screen. The probe items were identical to the memory items and their orientation deviated from the test item orientation by ± 13°, 19°, 26°, 33°, or 40°^3^. A black fixation dot was presented at the center of the screen with 0.73° of visual angle during the delay intervals.

### Procedure

Prior to the experiment, participants completed training trials that were identical to the experimental trials. The experimental session consisted of 1,440 trials divided into 18 blocks. Between blocks, participants took self-paced breaks. After every four blocks, participants were asked to take a longer break. During the blocks, trials were presented automatically one after the other until the block ended.

The experimental task is illustrated in Figure 1A. Each trial began with an interval randomly jittered between 1,100 and 1,300 ms during which a fixation dot was displayed. The fixation dot was presented during each delay interval throughout the experiment. A memory item was presented for 250 ms, followed by a 900 ms delay interval. Next, the rotation instruction was displayed for 200 ms during which the participants were instructed to rotate the first memory item. The participants were informed that after rotation, the original first memory item would no longer be task relevant. This was followed by a delay interval sequence which consisted of a 900 ms pre-impulse delay, a 100 ms impulse display, followed by a 500 ms post-impulse delay. Next, an additional memory item was presented for 250 ms and was followed by another delay interval sequence. Next, a retrocue display was presented for 200 ms, which cued the target item: indicating “1” for the rotation product, or “2” for the additional memory item. This was followed by another delay interval sequence. Finally, the probe orientation was displayed for 200 ms and was followed by an additional 1,100 ms response interval that contained a fixation dot. During this, the participants were asked to respond whether the probe orientation was rotated CW or CCW compared to the target item orientation, by pressing the A or L letter keys respectively. As feedback, the fixation dot turned green (RGB = [0, 255, 0]) for correct or red (RGB = [255, 0, 0]) for incorrect responses, for 150 ms immediately after response.

### EEG acquisition and preprocessing

EEG from 64 Ag/AgCI passive electrodes (Easycap Gmbh, Herrsching, Germany), positioned in accordance with the 10-5 system was recorded via an NeurOne Tesla AC amplifier (Bittium Biosignals Ltd, Kuopio, Finland) at 1000 Hz. Recording was done with a 250 Hz low-pass filter. The online reference electrode was FCz and the ground electrode was AFz. Four external electrodes were placed at the outer canthi and above and below the right eye. Their signal was recorded as bipolar horizontal and vertical EOG channels, but these were not analyzed further. All electrode impedances were kept below 10 kΩ at the beginning of the experiment, and checked again halfway during the experiment, after nine blocks.

Preprocessing of the EEG data was carried out using the EEGLAB (2024.2; Delorme & Makeig, 2004) toolbox in MATLAB (R2024a). Data was downsampled to 500 Hz and band-pass filtered (0.1 – 40 Hz). Noisy channels were visually identified and replaced using spherical interpolation. The data were re-referenced to the global average of all electrodes. Data were separately epoched for impulse 1, impulse 2 and impulse 3 between −100 ms before and 600 ms after impulse onset. Eye blink artefacts were labeled and removed from the data using independent component analysis (ICA). The data was then converted into the FieldTrip format (Oostenveld et al., 2011). Epochs with artifacts were excluded using the ‘ft_reject_visual.m’ summary method, if they met one of the following criteria: (1) signal variance within any channel exceeding 1500 µV, (2) absolute z-values exceeding 6 where the mean and standard deviation were computed over all time points and epochs per channel (3) absolute amplitudes exceeding 120 µV within any channel. We had predefined an exclusion criterion that a participant’s data would be excluded from the EEG and behavioral analyses if more than 30% of the impulse 1, impulse 2 or impulse 3 epochs were marked for exclusion. However, none of the participants met this criterion.

Subsequent EEG analyses were performed from the 17 parietal-occipital electrodes, as also used in previous WM decoding studies (Ataseven et al., 2026; Wolff et al., 2017; P7, P5, P3, P1, Pz, P2, P4, P6, P8, PO7, PO3, POz, PO4, PO8, O1, Oz, O2).

### Time-course decoding

The original and the rotation product of the first item were decoded from all three impulse epochs. The additional item was decoded from impulse 2 and impulse 3. The data was dynamically baselined, by centering each time point over a 100 ms time window, averaging over this time window per channel and epoch, and finally subtracting this average from all samples within the time window. Epochs were divided into eight cross validation folds where the number of trials in all folds per orientation was equalized via random sampling. Each fold was used as the test set once while the remaining folds were used as the training set. The training set was used to compute the covariance matrix across trials, and multivariate averages across trials for each orientation and each time point. In the test set, Mahalanobis distances were computed between each trial and these averages. For each orientation, the computed distances were averaged, sign-reversed and mean-centered. Finally, the resultant distances were cosine-weighted. The result was a decoding score that quantified the amplitude of the cosine-shaped tuning curve per time point. To avoid a selection bias while equalizing the number of trials per fold, this procedure was repeated across 100 repetitions and the average of the repetitions was used as the decoding score.

### Significance testing of the decoding

Average decoding accuracies were computed in the 0-300 ms interval following each impulse onset^4^. These were tested against zero with a Bayesian one-sample t-test, separately for the original item, rotation product for impulse 1, and the additional item, which was included in the tests for impulse 2 and impulse 3 (i.e., after it had been shown). For impulse 3, the test was done separately for trials in which the rotation product and the additional item was cued. BF_10_ < 0.333 or BF_10_ > 3 were set as thresholds for substantial evidence for or against above-chance decoding, following Wetzels and Wagenmakers (2012). Note that since each hypothesis in our preregistration makes a specific prediction for the three analysis windows, the combined Bayesian evidence for either hypothesis would effectively be larger than the BFs from the independent tests.

In the time-course analyses, non-parametric cluster-based permutation tests against a null distribution were used to test for significant decoding. First, the null distribution was generated by randomly flipping the sign of the trial-averaged time-course decoding scores from each participant over 10,000 simulations and recomputing the mean at each time point. Then, a cluster-corrected test compared the trial-averaged time-course decoding scores per participant against this null distribution to mark statistically significant clusters (p < 0.05). These clusters were defined by first grouping the neighboring time points where the mean exceeded the permutation-quantile threshold (0.05) and scoring the groups with the sum of their samplewise amplitudes.

### Analysis of behavioral data

We base the majority of our behavioral analyses on hierarchical binomial regression models predicting CW responses, using a cauchy link function that consistently outperformed probit- and logistic regression (Ataseven et al., 2026). Per specific analysis, we first use the AIC of fitted models to determine which combination of predictors of interest defined a ‘best’ model. In the main text, we quantify evidence for specific predictors by omitting them from the best model, then determining the p-value via chi-square likelihood ratio tests. In cases where we show absence of evidence for a specific predictor term, we add it to the best model and compare the resulting models. All models of behavior included fixed effects of probe distance with respect to the probed item, as well as a random intercept and a random slope term for probe distance. Every time we identified a best model, we also determined support for including additional random slopes, but these were not supported. Note that in models like this, effects on behavioral precision are characterized by a modulation of the slope coefficient of the psychometric curve, which is defined as an interaction with probe distance. Therefore, most model comparisons involve two degrees of freedom.

For the initial assessment of behavior, we investigated the effects of probe distance, retrocue (additional item cued or rotation product cued), and whether any mental rotation took place during the trial (yes or no). Based on AIC, the best model included all these terms and their two- and three-way interactions, which is inconsistent with results from likelihood ratio tests (reported in the results section), revealing that support for the inclusion of the three-way interaction and the two-way interaction between retrocue and any rotation was not significant. Therefore, we treated this simpler model as the ‘best’ model, and significance tests reported in the main text are derived from comparisons to this simpler model.

To assess the effects of mental rotation we separately analyzed trials with different retrocues. For both trial types, we evaluated the effect of having any mental rotation at all (yes, no), and the magnitude of this mental rotation (30°, 60°, 90°) coded as a linear predictor; specifically assessing their interaction with probe distance. As indicated in the results, we found that the best model for both data subsets included a term for the effect of any mental rotation, but only on trials where the rotation product was cued, a term for the magnitude of rotation was included in the best model. To visualize these precision effects in an intuitive manner, we fit nonhierarchical versions of this model, including terms for any rotation and rotation magnitude, independently to the data from individual participants (Figure 1B). Using estimated marginal means, we present the mean accuracies and 95% within-subject confidence intervals (Cousineau, 2005; Morey, 2008) across conditions as the insets of the figure.

To determine whether the responses to cued items were biased by the other two items, we again split the trials between different retrocue types, and discarded trials without mental rotation, as the rotation product is undefined on those trials. Next, we determined three probe distances per trial: to the rotation product, the original item, and the additional item. The models testing for bias effects initially included three fixed effects terms: for the distance to the cued item, as well as terms for the distance to the other two items. We quantified the evidence for either bias term by iteratively omitting them from the model. To visualize these resulting bias effects in our data more intuitively, we again turned to estimated marginal means from models fit to individual participants. Per participant, we used the model including all terms to predict the percentage increase in CW responses given a 10-degree increase in CW probe distance to either other item (see Figure 1C).

### Trialwise regression on decoding strength and behavior

We assessed the correlations between trial-wise decoding scores z-scored per participant using only trials where the rotation product was cued and mental rotation took place. In these trials, we tested different linear mixed-effects models predicting the decoding score of the rotation product, as a function of the decoding score of the original item and of the additional item. The resulting best model included a fixed effect for the original item decoding, as well as a correlated random slope term for this predictor. The coefficients across participants for this model showed that the resulting positive relation was highly consistent across participants (as depicted in Figure 4A).

We subsequently assessed whether either decoding score affected behavioral precision by taking the previously best behavioral model for these trials as identified in the previous section: This model included predictor terms for probe distance interacting with rotation magnitude, and a bias term for the probe distance with respect to the uncued additional item. Subsequently, we added predictor terms reflecting the decoding scores for each of the items, z-scored per participant, to test whether either of these terms would exert an additional effect on behavior, and found that both the original item and rotation product decoding scores interacted with probe distance, indicating a modulation of precision. We subsequently visualized this modulation of precision by taking this best behavioral model, fitting it separately to each participants’ data, and predicting accuracy at three levels of decoding (Figure 4B, corresponding to the insets of Figure 2A, but evaluated at three levels of decoding cf. the bins in Figure 4B).

## Acknowledgement

We thank Rama Alnabusy for her efforts in data collection.

## Author Contributions

Conceptualisation: NA, ŞÖ, EGA, DS, WK

Data curation: NA, ŞÖ

Formal analysis: NA, WK

Funding acquisition: No external funding

Investigation: NA, ŞÖ, EGA, DS, WK

Methodology: NA, ŞÖ, EGA, DS, WK

Project administration: NA

Resources: DS Software: NA, WK

Supervision: EGA, DS, WK

Validation: NA, WK

Visualization: NA, WK

Writing - original draft: NA

Writing - review & editing: NA, ŞÖ, EGA, DS, WK

## Footnotes

1 Time-course decoding uses features from a 100 ms sliding window. Decodability might therefore come to expression earlier than in the raw data. We present ERPs (Supplementary Figure 1) time-locked to each impulse display to confirm that the genuine visual response aligns with the recorded visual onset at time 0.

2 In our preregistration, we specified a Bayesian stopping rule according to which we would stop data collection whenever the hypothesis tests crossed the specified threshold in either direction, or if the maximum of 40 participants were recruited. Due to time constraints, the analysis was not run after each incoming dataset, and all 40 participants were recruited and all data were further analyzed.

3 For the first four participants, the probe deviated from the target item by 9°, 15°, 22°, 30°, and 39°. We observed these distances made the task rather difficult, and used larger probe distances for the remaining participants.

4 In the preregistration, this interval was specified as 100-400 ms following each impulse onset as prior studies (Kandemir & Akyurek, 2023; Wolff et al., 2020) have observed that decoding of the orientations were onsetted from 100 ms after stimulus. However, in our study, the decoding was observed earlier, starting shortly after the impulse onset. Therefore, we deviated from the preregistered interval in order to best represent the results for the data at hand.

