## Supplementary Material for "Mentally transformed representations in memory are linked to their originals"

Supplementary Materials

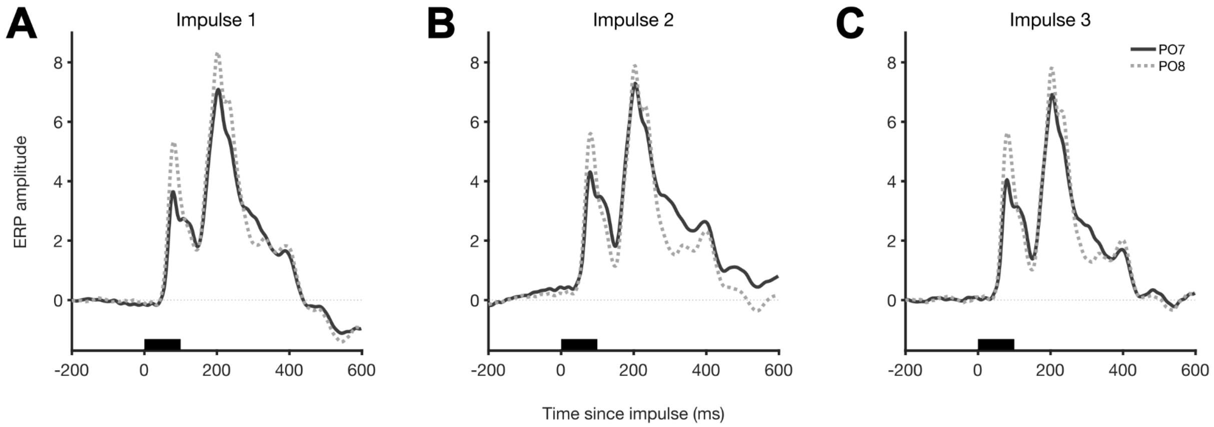


**Figure 1S. Event-related potentials during delay intervals.** The time-course plots are time-locked to the impulse onset in each delay marked by the black lines over the x-axes. The lines show the mean amplitude from electrodes PO7 (solid) and PO8 (dashed).
